# Brain regions with positive and negative task-evoked responses engage in cooperative rather than competitive interactions

**DOI:** 10.1101/687897

**Authors:** Kelsey D. Csumitta, Alexandra Ossowski, Adrian W. Gilmore, Stephen J. Gotts, Alex Martin

## Abstract

Brain regions displaying task-induced decreases in activity, sometimes referred to as “Task-Negative”, have been proposed to reflect cortico-cortical competition from regions showing task-induced increases in activity, sometimes referred to as “Task-Positive”. If these functional networks are competitive, trial-level fMRI BOLD responses across voxels exhibiting Positive and Negative responses should be anti-correlated. Additionally, the correlation between the BOLD response and behavior in Positive and Negative regions should have opposite sign. Here, we test these predictions using trial-to-trial variability in an object naming task. In contrast to the negative coupling proposed by a competition model, we find positive coupling between Positive and Negative regions and show that this coupling aligns with trial-level behavioral correlations. We provide evidence that these cooperative interactions are not due to a non-specific global factor and discuss alternative proposals, such as thalamo-cortical gating.

**Highlights:**

- Tested competitive interactions among positive and negative task-evoked regions
- Positive and negative regions exhibited positive coupling at trial and run levels
- Smaller competitive effects seen in BOLD-behavior correlations
- Strong cooperative effects not due to a non-specific common global factor

**eTOC:** Csumitta et al. show that positive and negative BOLD responses do not exhibit cortical competition among corresponding brain regions. Rather, these networks display positive coupling. The spatial distribution of positive coupling aligned with behavioral correlations, suggesting normal, cooperative network interactions.

## Introduction

Recent work in the fields of cognitive and systems neuroscience has been dedicated to understanding the dynamics of large-scale and fine-grained brain networks (Doucet et al., 2011; Gratton et al., 2018; Petersen and Sporns, 2015; Power et al., 2014; Shirer et al., 2012; Yeo et al., 2011). While engaged in a cognitive task during a BOLD fMRI experiment, such as making decisions about visual stimuli, brain regions thought to be most involved in the visual and decision demands of the task typically exhibit a positive, above-baseline BOLD response, often referred to as “Task-Positive” (Fox et al., 2005b) (referred to hereafter as “Positive” responses). At the same time, other regions (commonly in limbic-related temporal, medial frontal, medial parietal, and lateral parietal cortices) generally exhibit a transient negative, below-baseline BOLD response (Shulman et al., 1997), often referred to as “Task-Negative” (Fox et al., 2005b) (referred to hereafter as “Negative” responses). Negative responses have been suggested to reflect a variety of mechanisms and cognitive states, such as deviation from a “default mode” of processing (Raichle et al., 2001), a shift from internal to external attentional states (Andrews-Hanna et al., 2010a; Golland et al., 2007; Preminger et al., 2011; Simpson et al., 2001), and memory retrieval and memory-based scene construction (Andrews-Hanna et al., 2010b; Sestieri et al., 2011). In the absence of an overt task, Positive and Negative regions identified in resting-state brain activity have been argued to display anti-correlated temporal interaction (Fox et al., 2005b; Fox et al., 2009). The presence of opposing responses during a wide range of tasks, as well as during resting brain activity, may reflect the presence of stable, inherently competitive brain dynamics across these neuroanatomical systems.

However, two main challenges have been raised for the competition hypothesis. The first challenge is that regions can change from Negative to Positive depending on experimental context, undermining the view that stable anti-correlated brain circuits are consistent between task and rest. For example, Negative responses are typically found in medial prefrontal cortex, posterior cingulate, superior temporal sulcus (STS), and the temporal pole while making decisions about pictured objects. However, Positive responses are found in these same regions when the task involves comprehension of verbal stimuli (Binder et al., 2010; Ezra et al., 2010; Mar, 2011), learning facts verbally about unfamiliar people (Simmons et al., 2010), or interpreting social content (Molenberghs et al., 2016; Spreng and Grady, 2010; Wheatley et al., 2007). Similarly, “Negative” and “Positive” regions have been shown to reverse their responses within the same subjects when task conditions switch from requiring visuospatial attention to memory-based processes (Sestieri et al., 2010; Spreng et al., 2010). These findings highlight the context-specific nature of Positive and Negative responses. A second main challenge for the competition view is that anti-correlated dynamics between Positive and Negative regions at rest depend partly on methodological factors. Murphy and colleagues revealed that regressing the whole-brain or “global” signal during preprocessing can lead positive correlations or those near zero to become negative after the regression (Anderson et al., 2011; Murphy et al., 2009; Saad et al., 2012; Weissenbacher et al., 2009).

Taken together, these relationships and methodological factors motivated us to re-examine the mechanistic proposal in (Fox et al., 2005b) and to do so in the context of a specific experimental task rather than during the resting state: In a context in which both Positive and Negative responses are observed, are these responses really generated by competitive interactions between networks? In principle, interactions could be competitive, cooperative, or even independent, with regions dynamically coupling and uncoupling depending on the momentary task demands (see Figure 1).

**Figure 1.**
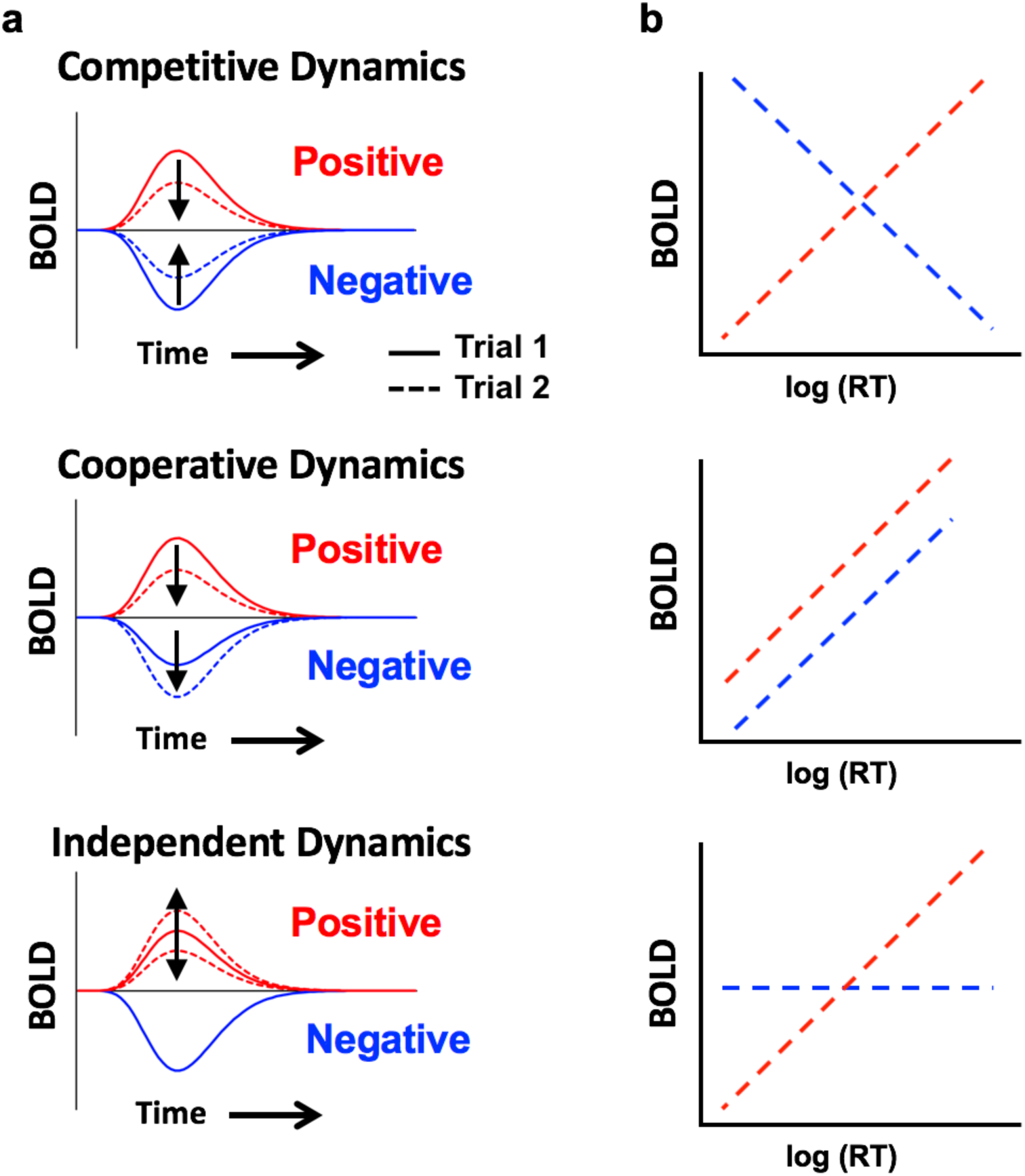
Competitive, cooperative, and independent dynamics predictions. **(a)** In line with the competition proposal (top), increases in the BOLD signal in one network are coupled with a decrease in BOLD in the opposing network. In contrast, under cooperative dynamics (middle) a decrease in BOLD in one network would lead to a concurrent decrease in the other network. Under independence (bottom), fluctuations would have no consistent relationship between the two types of regions, such as when changes in one region are not present in the other. **(b)** With competition, the correlation between trial-level BOLD response amplitudes and corresponding behavioral performance should be of opposing sign in Positive and Negative regions (top). Under cooperative dynamics, BOLD-behavioral correlations should have the same sign in Positive and Negative regions (middle). Independent dynamics predict that the slopes should be unrelated, perhaps with non-zero slope in Positive regions and zero slope in Negative regions.

In the current study, we use BOLD fMRI to examine what type of dynamics accompany Positive and Negative responses during visual object identification. We employ a slow event-related design in order to isolate the responses to individual trials, permitting independent calculation of trial-level response amplitude variability from the shape and sign of the trial-averaged evoked responses (Positive versus Negative). In the context of this design, we can formulate the following two predictions:

1. If competitive dynamics explain Positive and Negative responses (i.e. they reflect a single underlying mechanism), then the amplitudes of the corresponding trial-level responses should exhibit negative correlations over the full spatial extent of these regions (see Figure 1a). In contrast, positive correlations should be observed with cooperative dynamics and correlations near zero should be observed for independence.

2. Further, if inputs from Positive to Negative regions (or vice-versa) are effectively inhibitory or suppressive, then any local correlation between the trial-level BOLD response amplitude and corresponding behavioral performance (in this case, RT) (Rao et al., 2014; Yarkoni et al., 2009) in Positive regions should be reflected in Negative regions with a reversed slope (see Figure 1b). Under cooperative dynamics, both regions with Positive and Negative responses should exhibit non-zero slopes with the same slope direction (either both positive or both negative). Under independence, we would expect a non-zero slope for regions with Positive responses and perhaps zero slope for regions with Negative responses (corresponding to a lack of relationship between the two types of regions).

Both of these predictions were assessed using BOLD fMRI in 40 participants, who were scanned while naming aloud pictures of common objects. Naming responses and corresponding RTs were recorded using an MRI-compatible microphone. We employed multi-echo independent component analysis (ME-ICA) to preprocess the data (Kundu et al., 2012; Kundu et al., 2017), which differentiates BOLD from non-BOLD components based on the decay properties of the BOLD signal across multiple read-outs (TEs) during each TR, allowing an automated means of removing focal non-BOLD artifacts such as are produced by motion (Power et al., 2018). Following the assessment of the main predictions, we also examined the potential for residual global sources of variation (artifactual or otherwise) to mediate the observed results.

## Results

### Identification of Positive and Negative regions

Participants completed a picture naming task consisting of 100 unique photographic images of animals, plants, and everyday objects presented across 2 scan runs, each consisting of 50 images. We employed a slow event-related fMRI design to permit the isolation of the peak BOLD response associated with individual trials. Responses and corresponding naming RTs were obtained using a noise-cancelling microphone for each trial. Positive and Negative voxels in the brain were identified during picture naming using the group-average beta coefficients. Group-average betas for each run were tested against baseline using one-sample t-tests to determine if voxels were Positive (exhibiting a positive t-statistic) or Negative (exhibiting a negative t-statistic), with correction for whole-brain comparisons performed using False Discovery Rate (FDR, q<.05; (Genovese et al., 2002)). We then required replication across the two runs (either significantly Positive or Negative in both runs) in order to define Positive and Negative regions of interest (ROIs). At the FDR-corrected statistical threshold, Positive voxels constituted one large mask of contiguous voxels (Figure 2a; Table S1), whereas Negative voxels were organized into eight more restricted Negative ROIs (see Figure 2; Table S1). The 8 Negative ROIs were then used as seeds for a form of whole-brain functional connectivity analysis, permitting us to examine whether the trial-level co-fluctuations in response amplitude were positive or negative with the voxels in the Positive mask.

**Figure 2.**
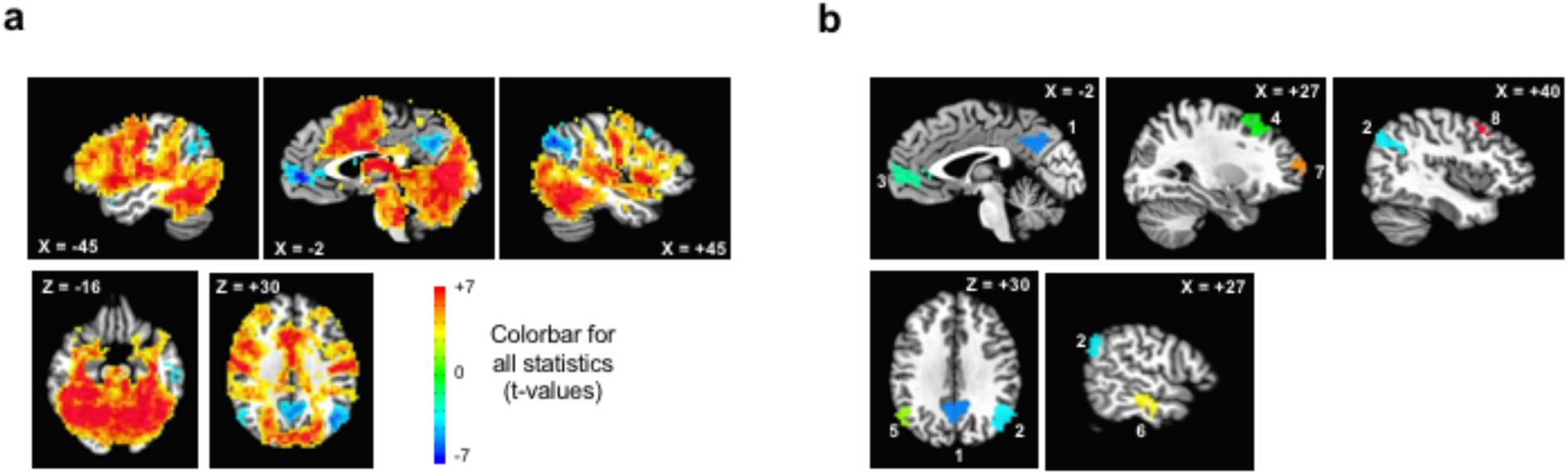
Positive and Negative regions. **(a)** Positive voxels (red) vs. Negative voxels (blue) during object naming (p<0.05, q<0.05 FDR). **(b)** Negative ROIs replicating across runs (p<0.05, q<0.05 FDR). See also Table S1.

### Prediction 1 results: correlations of trial-level responses

If Positive and Negative responses reflect stable cortico-cortical competition, then the amplitudes of the corresponding trial-level responses should exhibit negative correlations over the full spatial extent of these regions. In other words, if a large Positive response is observed to a particular trial in Positive voxels, competition predicts a larger Negative response in Negative voxels. To test this prediction, we first isolated the peak BOLD responses for each correct trial from the de-noised data (an average of the 2nd and 3rd TRs on each trial, chosen to be consistent with a typical peak BOLD response of 4-8 s post-stimulus). We then performed a whole-brain, seed-based correlation analysis from the 8 Negative ROIs using these single-trial responses as “item series” (rather than time series). In contrast to the competitive dynamics proposal, there were no significant (or even numerical) anti-correlations found with any Positive voxels at the group level (see Figure 3a). Indeed, these correlations were modestly positive, ranging from 0.2 to 0.3 in magnitude (see Table S2).

**Figure 3.**
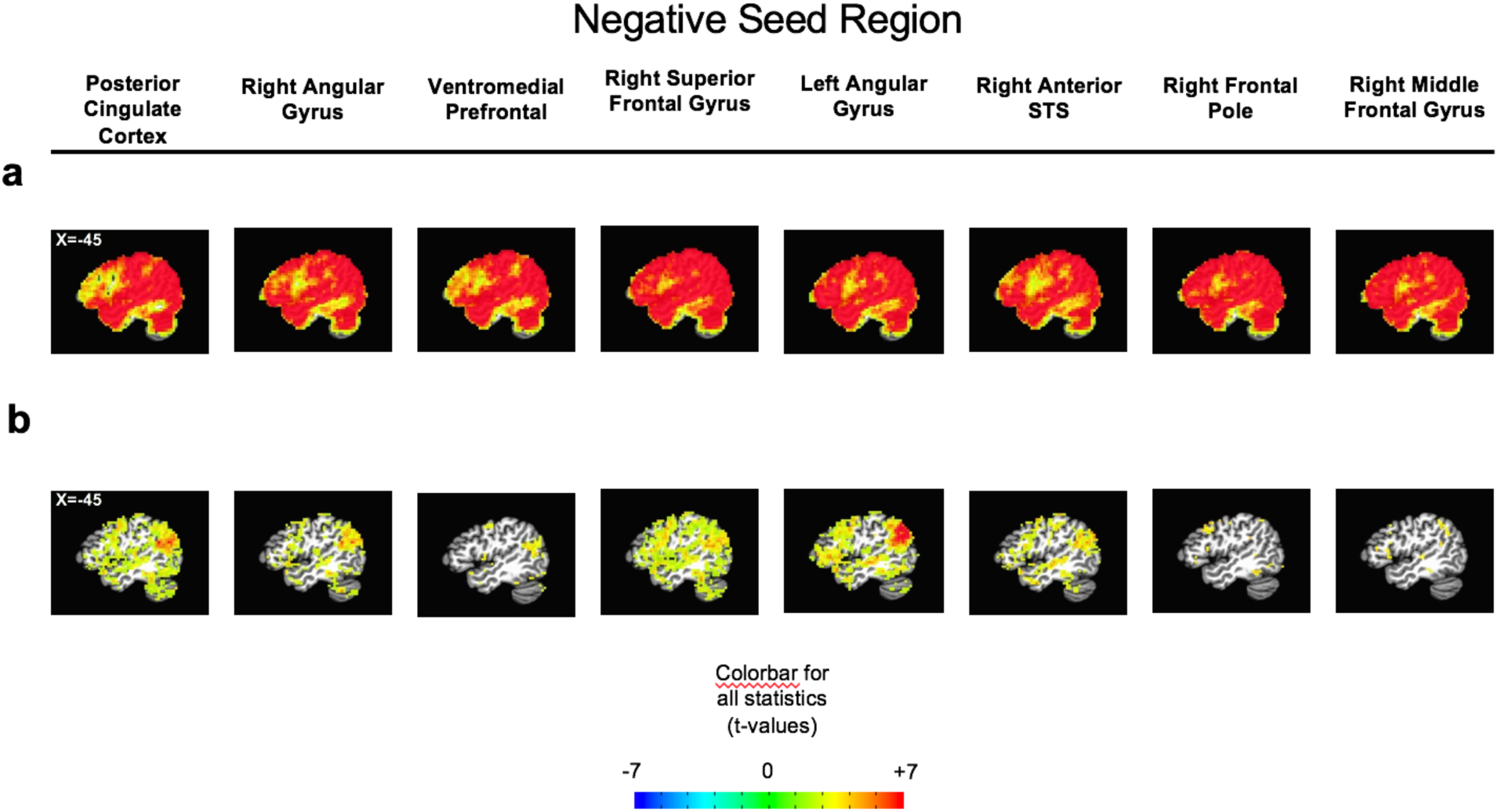
Prediction 1 results: seed-based correlation analyses. **(a)** Whole-brain correlations using the 8 Negative ROIs as seeds (p<0.05, q<0.05 FDR), partialling trial-level artifact measures. **(b)** Beta coefficient covariation analysis using the change from run 1 to run 2 (p<0.05, q<0.05 FDR), partialling run-level artifact measures. See also Table S2.

#### Contribution of residual artifacts

It is well-known that head motion and other factors (e.g. respiration) can continue to impact measures of functional connectivity after de-noising. The expected impact would be to add modest positive correlations to such measures, potentially consistent with the current pattern of results (Power et al., 2017). The ME-ICA denoising approach employed here has previously been shown to remove pure effects of head motion well, while failing to eliminate slower effects of respiration (Power et al., 2018). We therefore evaluated whether trial-level motion and other artifact sources were responsible for the positive correlations found between Negative ROIs and the Positive voxels. Trial-level estimates of motion were constructed from Framewise Displacement measures, as were two more omnibus measures of artifact magnitude based on temporal signal-to-noise ratio (see Methods for details). These trial-level estimates were then partialled from the trial-level correlations between Negative and Positive voxels, with the partial correlations tested against zero. The partial correlations between Negative and Positive were modestly positive (see Figure 3a).

#### Covariation of beta coefficients across runs

The trial-level amplitudes used in the previous analyses were not based on a multiple regression model but were instead simply “notched” out of the de-noised time series during the period of the expected peak BOLD response. A useful property of beta coefficients in multiple regression is that, while they are influenced by noise in the independent variables (Frost and Thompson, 2000; Spearman, 1904) they do not depend on noise in the dependent variable (in this case, BOLD data). One might therefore expect covariation among beta coefficients in Positive and Negative regions to be an important alternative metric related to the competition hypothesis (see also (Rissman et al., 2004)). The beta coefficients from the task regression that were calculated on the separate runs (runs 1 and 2) were used for this analysis. The change in beta coefficient between run 1 and run 2 in Negative ROIs were correlated across participants with changes in their respective Positive voxels. As with the trial-level analyses, the competition hypothesis predicts that an increase (decrease) in betas from run 1 to run 2 in Negative ROIs should be accompanied by a decrease (increase) in betas in Positive voxels. These analyses also covaried run-level changes in head motion and temporal signal to noise ratio (see Methods for details). As with the previous analyses, changes in run-level beta coefficients showed positive correlations between Negative ROIs and Positive voxels (see Figure 3b). Indeed, the positive correlations were present across all three of these analyses (in the 0.2-0.3 range; Table S2). Taken together, these results are in conflict with the competition hypothesis.

### Prediction 2 results: correlation between trial-level BOLD magnitude and behavior

In the previous analyses related to prediction 1, we failed to find evidence of competition between Positive and Negative brain regions at the trial and run levels. Instead, we observed modest positive coupling consistent with cooperative dynamics, and this coupling did not obviously relate to artifact measures such as head motion or temporal signal-to-noise ratio. The competitive dynamics proposal also makes a second prediction for behavioral correlations: if Positive BOLD responses are positively correlated with RT, Negative BOLD responses should be negatively correlated with RT (Rao et al., 2014; Yarkoni et al., 2009). To test this prediction, we first examined the correlation of the BOLD response magnitude and corresponding RT for each correct trial, per participant. As in the correlation analyses for prediction 1, we then partialled trial-level estimates of residual artifacts and repeated the analysis.

The correlation between the trial-level BOLD magnitude and RT was positive overall in Positive voxels but negative in 6 out of 8 Negative ROIs (see Figures 4a and 4b, Table S3). In the remaining two Negative ROIs, the correlations were near zero. Taken in isolation, these results support the competition hypothesis. However, the mean correlation values overall were much smaller than the positive coupling levels observed between Negative ROIs and Positive voxels (values less than 0.1 compared to values between 0.2 and 0.3; see Table S3), and negative BOLD-RT correlations were also observed in several Positive regions (see Figure 4c), particularly in the supramarginal and inferior frontal gyri, bilaterally, as well as in the right putamen. The mixing of positive and negative BOLD-RT slopes in different collections of Positive voxels would appear to undermine a single unitary cooperative or competitive explanation. Instead, these results seem to support competition between Positive and Negative regions in this task for some portion of the overall trial-level variance (and perhaps amongst different collections of Positive voxels).

**Figure 4.**
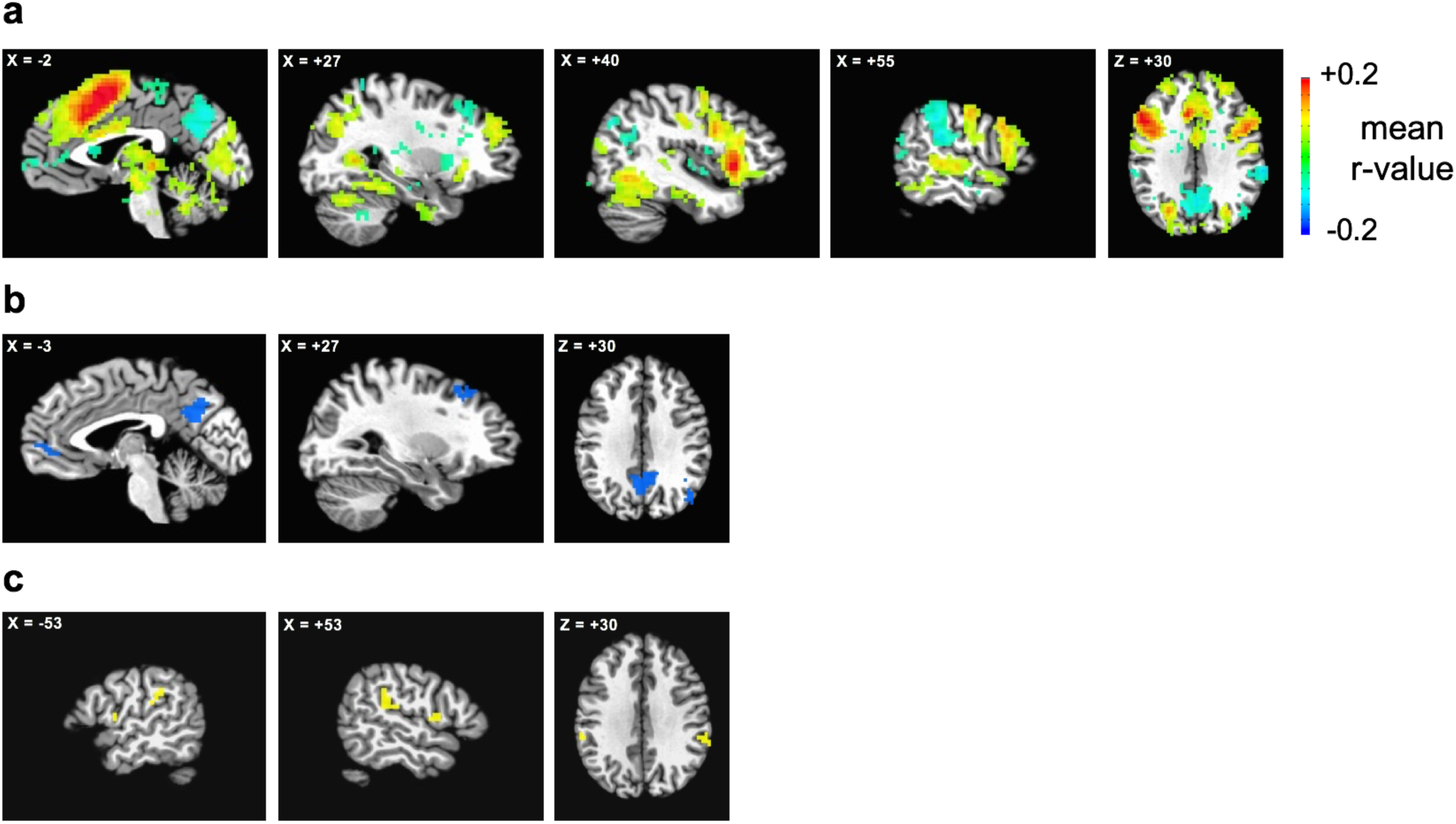
Prediction 2 results: voxelwise correlation between trial-level BOLD magnitude and RT. **(a)** Mean correlation between voxelwise BOLD magnitude and RT, regressing nuisance covariates (item-specific FD and two models of SD, p<0.05, q<0.05 (FDR) for all), with no additional masking. **(b) 6 of 8** regions that are Negative show a negative correlation with RT, **potentially** in line with a competitive dynamics proposal. **(c) A subset of Positive** voxels also show a negative correlation with RT, **which was unanticipated and is difficult to account for using a competitive dynamics proposal**. See also Table S3.

Given the seeming conflict between the results related to predictions 1 and 2, we next examined to what extent the negative coupling between Positive and Negative voxels in behavior (prediction 2) contributes to the overall estimates of positive coupling found in prediction 1. To evaluate this, we calculated the seed-based correlation between each Negative ROI and all of the Positive voxels (independent of the nuisance variables), transforming the correlations to *R*^*2*^ in terms of the original trial-wise amplitude variance and then averaging across all voxels in the Positive mask. Next, we recalculated these values after removing the relationships with RT. Removing RT had little or no additional effect on the level of coupling when averaging over the entire Positive mask (Figure 5). The *R*^*2*^ of trial-level variance explained by RT in the local timeseries of the Negative ROIs and Positive Voxels is shown in Figure 5 for reference, with values much smaller than the level of positive coupling indicated by the *R*^*2*^ values shared between Negative and Positive voxels when excluding RT. These results indicate that while competition may be present in a portion of variance, this portion is relatively small, with the majority of coupling between Positive and Negative voxels being positive, consistent more with cooperative dynamics (as in Figure 1a).

**Figure 5.**
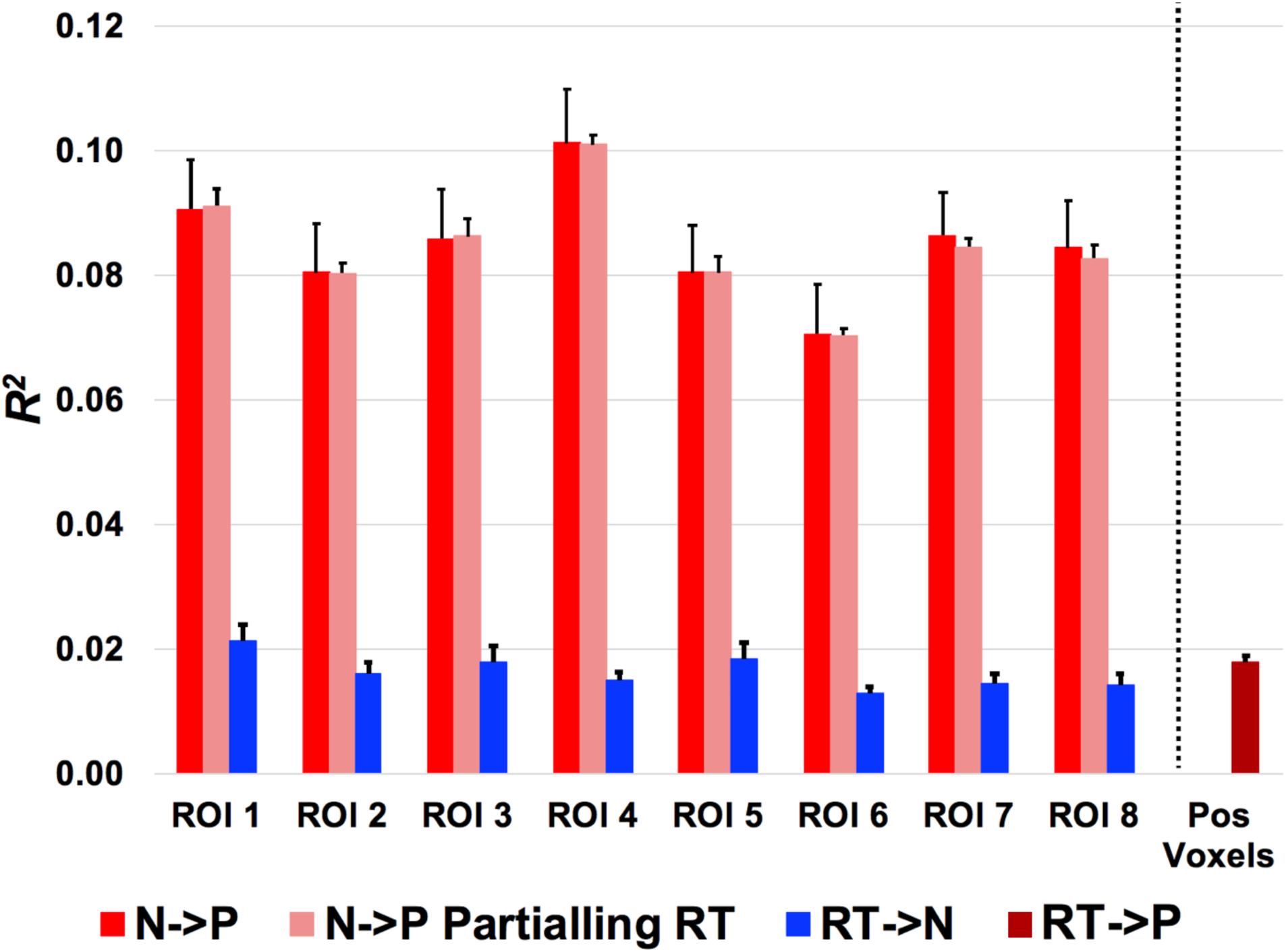
Positive coupling between Negative and Positive voxels is stronger than RT-related variance. Variance in the trial-level responses that is coupled between Negative ROIs and Positive voxels (N->P, independent variable N and dependent variable P) is shown in terms of *R*^*2*^ after removing nuisance variables. These estimates show little or no change after removing the putative “competition-related” variance related to RT (N->P Partialling RT). Local trial-level variance in Negative ROIs (RT->N) and Positive voxels (RT->P) explained by RT is also small by comparison.

### Global contributions to Positive and Negative Regions

Thus far, our results have indicated predominantly positive coupling between Positive and Negative regions with some smaller competitive effects seen related to behavior (RT). In the examination of predictions 1 and 2 above, we took care to control for the contribution of global artifacts such as head motion, as well as any other source of variation reflected in average measures of temporal signal-to-noise ratio. However, an additional possibility is that other sources of variation with more neurogenic bases, such as arousal (e.g. (Chang et al., 2016); (Deco et al., 2014; Fox et al., 2005a; Jones, 2005; Massimini et al., 2005; Saper et al., 2005)), contribute joint variance to Positive and Negative regions, leading to positive trial-level correlations. Previous studies of arousal effects on functional connectivity have shown a distinct pattern of positive correlations amongst cortical regions and negative correlations with thalamus and cerebellum (e.g. (Chang et al., 2016; Feige et al., 2005; Goldman et al., 2002; Liu et al., 2012; Moosmann et al., 2003; Olbrich et al., 2009; Ong et al., 2015; Poudel et al., 2014)). If the positive correlations truly reflect competition plus a global common factor (such as arousal), we would expect that the maximum positive correlations from Negative regions onto Positive voxels would occur outside of those voxels most engaged in the performance of the task (with the minimum positive correlations occurring at the locations of maximum competition related to RT). We therefore examined the relative distribution of positive correlations between Negative regions and voxels in the Positive mask.

In contrast to the expectation of arousal and other global effects plus competition, the strongest positive correlations from Negative regions tended to be in the same Positive voxels that were most associated with task performance in picture naming (see Figure 6). A more continuous comparison of these distinct correlation maps within the large Positive mask also revealed significant positive (Spearman rank) correlations between the seed-based beta weight results and the RT correlations, as well as between seed-based trial-level correlations and RT correlations (Table 1; see Methods for details). Finally, strong positive seed-based correlations from Negative regions were observed in thalamus and cerebellum, unlike previous studies of arousal effects. These results all undermine alternative global explanations of the positive correlations between Positive and Negative regions.

**Table 1.** Strongest positive correlations from Negative ROIs onto Positive Voxels match those correlated with behavior. Within the large Positive mask, significant positive (Spearman rank) correlations were found between seed-based beta weight correlations and RT correlations, as well as between seed-based trial-level correlations and RT correlations. *P*-values for the correlations were determined through Monte Carlo simulations using random maps with a spatial auto-correlation function matched to the original data. All significant *p*-values (*p*<.05) were also FDR-corrected (*q*<.05). No significantly negative correlations were found for either analysis. These results argue against global origins of positive coupling between Positive and Negative regions. Related to Figure 6.

| Negative ROIs | Seed-Based Beta<br>Coefficient Covariation<br>Analysis & RT Correlations |  | Seed-Based Trial-Level<br>Correlations & RT<br>Correlations |  |
| --- | --- | --- | --- | --- |
| | $r$ | $p$ | $r$ | $p$ |
| 1 Posterior Cingulate Cortex | 0.1507 | 0.0002 | 0.0205 | 0.6080 |
| 2 Right Angular Gyrus | 0.2498 | 0.0002 | 0.2073 | 0.0002 |
| 3 Ventromedial Prefrontal | -0.0473 | 0.1254 | 0.1055 | 0.0032 |
| 4 Right Superior Frontal Gyrus | 0.2554 | 0.0002 | 0.1993 | 0.0002 |
| 5 Left Angular Gyrus | 0.2431 | 0.0002 | 0.2670 | 0.0002 |
| 6 Right Anterior STS | 0.2386 | 0.0002 | 0.2465 | 0.0002 |
| 7 Right Frontal Pole | 0.2890 | 0.0002 | 0.2761 | 0.0002 |
| 8 Right Middle Frontal Gyrus | 0.3581 | 0.0002 | 0.3665 | 0.0002 |

**Figure 6.**
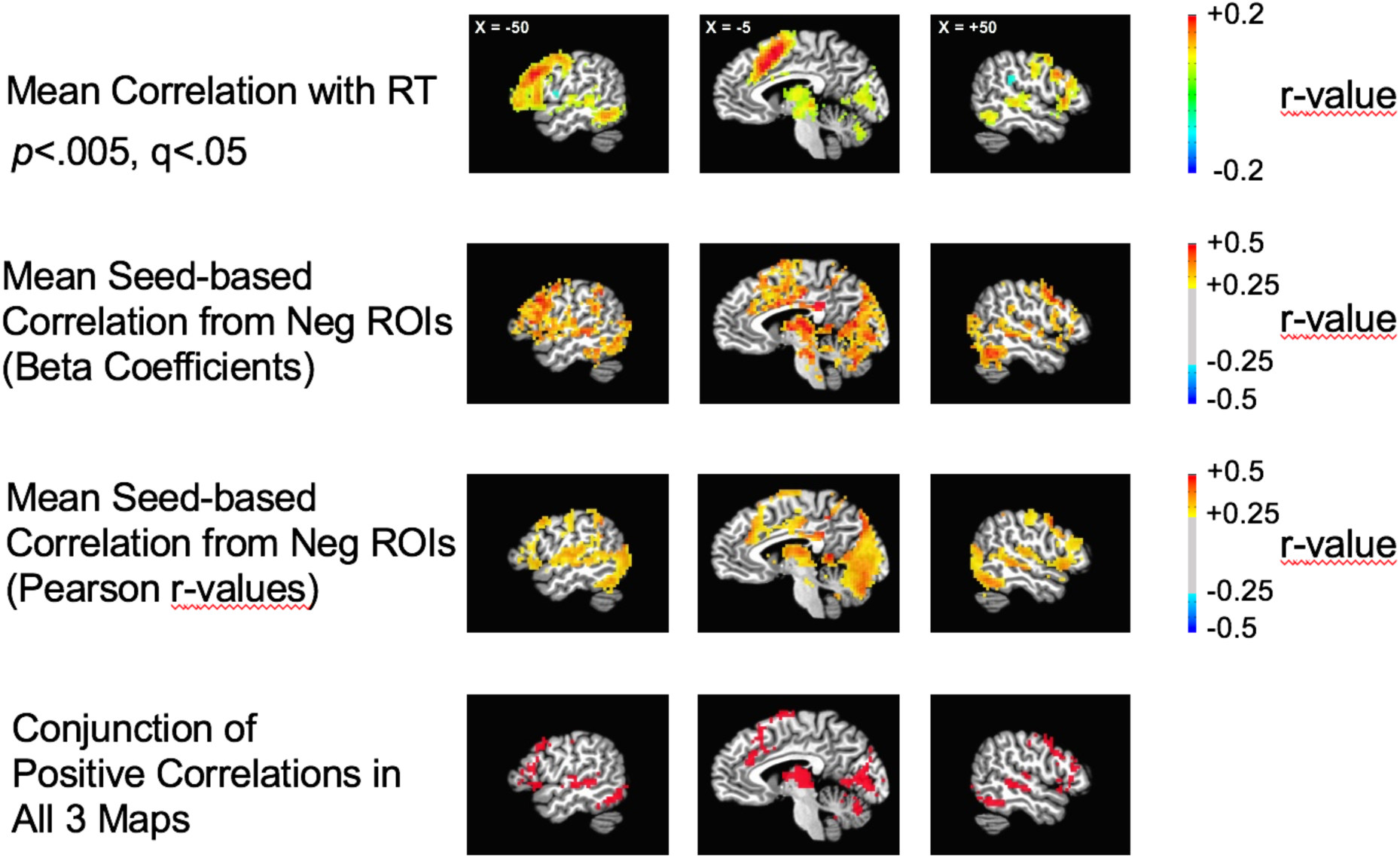
Strongest positive correlations from Negative ROIs onto Positive Voxels match those correlated with behavior. The average seed-based correlation from Negative (Neg) ROIs (averaged across all 8 Negative ROIs) onto Positive voxels tended to be most positive (>+.25) in the same voxels that were positively correlated with response times in picture naming (shown for reference in top row). This was true for both the beta coefficients, partialling nuisance measures (second row), as well as for item-series functional connectivity (Pearson r-values), also partialling nuisance measures (third row). Competition plus a non-specific common global factor (either nuisance or real, such as an arousal effect) would instead predict that the weakest (rather than strongest) positive correlations would be aligned with the RT correlations. The conjunction of positive correlations in the first three thresholded maps are shown in the bottom row.

## Discussion

In the current study, we tested the idea that Positive and Negative regions engage in competitive cortico-cortical interactions in the context of two predictions: 1) trial-level responses should be negatively correlated, and 2) correlations with behavior (RT) should exhibit a reversal of sign. Related to prediction 1, trial-level correlations were modestly positive rather than negative, and this positive sign had little apparent relation to the magnitude of residual artifacts such as head motion, nor did it resemble arousal or other global effects. Analyses of the change between runs in beta coefficients between runs, which are expected to depend less on motion and other artifacts, yielded convergent results. Related to prediction 2, some evidence of BOLD-RT slope reversal was found between Positive and Negative regions. However, these effects were small in magnitude, and when removed, the level of positive coupling observed was little affected.

Some comparable results have been previously reported in (Mayhew et al., 2016). Mayhew and colleagues conducted three separate stimulation experiments (visual, motor, and somatosensory) to induce Positive and Negative BOLD responses. At the single-trial level using the peak amplitudes from individual time-courses (as was done here), Positive and Negative BOLD responses were positively correlated. At the same time, group mean Negative and Positive response magnitudes were negatively correlated. This negative relation was due to concomitantly greater Positive and Negative mean responses at higher stimulus intensities, proposed to arise from stimulus intensity-dependent ipsilateral inhibition together with increased contralateral excitation (Klingner et al., 2010; Shmuel et al., 2002). Our study differs from this previous work in that our analyses of mean beta coefficients across participants (by run) indicate positive rather than negative correlations among Positive/Negative regions, and we further show that the spatial distribution of this positive coupling relates to BOLD-behavior correlations in the task, ruling out the contribution of non-specific “global” sources to the positive coupling.

Additional studies have found varying levels of positive coupling between Positive and Negative regions depending on experimental context. Fornito and colleagues investigated the competition proposal using a recollection task and found greater task-related positive coupling between Positive and Negative networks across participants was associated with more rapid memory recollection (Fornito et al., 2012). Further studies have found transient positive correlations between Positive and Negative systems during working memory tasks (Bluhm et al., 2011), task-selective recollection (Simons et al., 2008), and goal-based mental simulations (Gerlach et al., 2011; Spreng et al., 2010). The current study expands on the aforementioned work by examining the relationship of these phenomena to trial-level behavioral measures, examining the effects on beta coefficients as well as more standard functional connectivity measures, and thoroughly examining the contributions of more global sources of variation.

Overall, our results do not support dynamic competition between Positive and Negative regions and instead support cooperative functional interactions. Previous work used to establish dynamic competitive effects among Positive and Negative networks has included the global average brain signal as a nuisance regressor (e.g. (Fox et al., 2005b; Fransson, 2005; Greicius et al., 2003; Kelly et al., 2008)). Given that negative correlations are guaranteed for the lower half of the original correlation distribution, even when all correlations are initially positive (due to the re-centering of the full correlation distribution around 0; e.g. (Fox et al., 2009; Murphy et al., 2009; Saad et al., 2012)), one cannot cleanly evaluate the competition hypothesis under these conditions. The preprocessing in the current study utilized Multi-echo ICA (or ME-ICA, (Kundu et al., 2012)), which sorts BOLD from non-BOLD variation using the known decay properties of BOLD effects across multiple read-outs per TR. Unlike global signal regression, this form of preprocessing will retain relatively global sources of BOLD variation, as well as some residual global artifacts related to respiration (e.g. (Power et al., 2018)). Given that the positive correlations observed in the current study were unrelated to markers of these residual artifacts (e.g. Framewise Displacement, temporal signal-to-noise ratio) and that their spatial distribution aligned with the trial-level behavioral correlations (Figure 5), we would suggest that these correlations may relate in some manner to normal, cooperative network functioning and interaction. Furthermore, the results are comparable using trial-level variation and run-level variation of beta coefficients.

However, the presence of positive correlations among Positive and Negative regions presents a further puzzle. If Negative responses do not reflect competition from Positive regions, what do they reflect? Given the fact that regions can exhibit either Positive or Negative responses, depending on the choice of task, the mechanisms that generate the Negative responses are unlikely to reflect region-specific differences in the task-evoked BOLD response. One possibility that is consistent with current data is that Negative responses reflect a more centralized gating mechanism at the onset of each trial, perhaps mediated by structures such as the reticular nucleus of the thalamus (e.g. (Ferrarelli and Tononi, 2011; Guillery and Harting, 2003; Zikopoulos and Barbas, 2007). Cortico-cortical interactions that contribute to positive correlations at the trial-level could then have additive effects to the more centralized gating that determines the shape of the local evoked responses and that is apparently inverse between Negative and Positive regions. Future studies could examine this possibility by manipulating tasks using slow event-related designs within the same participants in such a way as to reverse Positive and Negative responses, tracking whether thalamic or other responses mediate this shift.

More generally, our current results highlight the utility of slow event-related designs for examining task-based functional connectivity. One perennial problem in task-based functional connectivity is that it is typically impossible to remove fully the evoked response in each voxel from the connectivity estimates. By including only the peak response period per trial, the shape of the evoked response has no influence on the connectivity estimates in the current study. In this context, connectivity instead reflects joint increases or decreases in the evoked response amplitudes in individual trials across pairs of voxels/regions. This approach can be fruitfully applied to many cognitive contexts that employ trial spacing of several seconds and that require interleaving of different trials types, permitting easy censoring of individual trials due to poor performance.

## Acknowledgements

We thank Shawn Milleville and Sarah E. Kalinowski for assisting with data collection and analysis. This study was supported by the National Institute of Mental Health, NIH, Division of Intramural Research (ZIAMH002920; ClinicalTrails.gov ID NCT00001360). The funders had no role in the study design, data collection and analysis, decision to publish, or preparation of the manuscript.

## Author Contributions

Conceived and designed the experiments: KDC, AO, AWG, SJG, AM Performed the experiments: AWG

Analyzed the data: KDC, AO, AWG, SJG, AM

Wrote the paper: KDC, AO, AWG, SJG, AM

## STAR Methods

### Lead Contact and Materials Availability

#### Lead Contact

Further information and requests for resources and reagents should be directed to and will be fulfilled by the Lead Contact, SJG.

#### Materials Availability Statement

This study did not generate new unique reagents.

### Experimental Model and Subject Details

#### Ethics Statement

Ethics approval for this study was granted by the NIH Institutional Review Board (protocol 93-M-0170, clinical trials number NCT00001360).

#### Participants

This study consisted of 40 participants (23 female, mean age: 24.5 years, range: 19 to 35) recruited from the NIH community and the DC metropolitan region. Participants were right-handed, neurologically healthy native English speakers with normal or corrected-to-normal vision. All participants granted informed consent and were monetarily compensated for their participation. Portions of these data have been reported previously in (Gilmore et al., 2019).

### Method Details

#### Experimental Stimuli

Participants completed a picture naming task consisting of 100 colored photographic images of animals, plants, and everyday objects. Images were presented across two lists (selected out of total of four, with two additional lists used in later parts of the experiment reported in (Gilmore et al., 2019)), with each list consisting of 50 unique images. Average lexical properties of picture names were equated across lists (omnibus *F* statistics all < 1) including lexical decision times on the names (mean RT = 632.3 ms, SD = 71.9 ms) and log HAL frequency determined by the English Lexicon Project database (mean = 8.57, SD = 1.54) (Balota et al., 2007). Additionally, average picture naming RT (mean = 826.8 ms, SD = 98.2 ms) was equated across lists based on pilot data on 60 additional participants (*F*(3, 196) = 0.736, *p* = .532).

Images were resized to 600 × 600 pixels and presented against a gray background (RGB value: 75, 75, 75) in the center of a 100 Hz MR-compatible monitor (screen resolution: 1920 × 1080 pixels) located at the head of the scanner bore using Presentation software (Neurobehavioral Systems). Trials began with a 1-s onset cue in the form of an orange fixation cross (48 point Arial type). Images were then presented for 300 ms and were followed by a white fixation cross for a variable period of 5.3 to 11.9 s.

#### Naming Task

Participants orally named 100 images presented across two 50-image scan runs. Participants were instructed to name each image aloud as quickly, accurately, and clearly as possible. Participants spoke into an MR-compatible microphone placed next to the head coil approximately 3-5 cm from the participant’s mouth. Accuracy for all participants across both runs each containing 50 pictures was approximately 90% correct. Only correct trials were included in our analyses. Median RT was approximately 960 ms (range: 697 s to 1270 ms) with a standard deviation of approximately 240 ms. These data were collected as part of a larger study on implicit and explicit memory effects (see (Gilmore et al., 2019)), and these two naming runs served as the pre-exposure phase, with all stimuli presented a single time with no repetitions. As such, the data were always collected prior to any other task conditions, and these later conditions are not discussed further here.

#### Recording Naming Responses during MRI

Spoken responses were captured with an Opto-Acoustics FOMRI-III NC MR-compatible microphone with built-in noise cancellation and routed into an M-Audio FastTrack Ultra 8-R USB audio interface. Responses were recorded with Adobe Audition. To calculate response RTs, the stimulus presentation computer emitted a square wave pulse at the onset of each trial and a custom Matlab program calculated the time difference between the square pulse onset and voice response for each trial.

#### MRI Methods

Images were acquired with a General Electric Discovery MR750 3.0T scanner using a 32-channel head coil. A high-resolution T1 structural image was obtained for each participant (TE = 3.47 ms, TR = 2.53 s, TI = 900 ms, flip angle = 7°, 172 slices with 1 ×1 ×1 mm voxels). Functional images were acquired using a BOLD-contrast sensitive multi-echo echo-planar sequence (Array Spatial Sensitivity Encoding Technique [ASSET] acceleration factor = 2, TEs = 12.5, 27.7, and 42.9 ms, TR = 2200 ms, flip angle = 75°, 64 ×64 matrix, in-plane resolution = 3.2 × 3.2 mm). Whole brain EPI volumes (MR frames) of 33 interleaved, 3.5-mm-thick oblique slices (manually aligned to the AC-PC axis) were obtained every 2.2 s.

Foam earplugs were worn by participants to attenuate scanner noise and participants’ head positions were stabilized using foam pillows. Each participant wore a respiration belt to monitor breathing, and heart rate recordings were obtained with a sensor placed on each participant’s left middle finger.

#### fMRI Data Preprocessing

Multi-echo functional images were preprocessed to attenuate noise and enable registration across participants using AFNI (Cox, 1996). Preprocessing steps included removing the first four volumes to allow T1 stabilization, despiking, slice-timing correction, and frame-by-frame rigid-body realignment to the first retained volume of each run. Data from all three acquired echoes were used to remove additional thermal and physiological noise with ME-ICA (Kundu et al., 2012; Kundu et al., 2017). This technique reduces thermal noise and separates BOLD from non-BOLD components (including motion and hardware artifacts) based on the known linear properties of the T_2_^*^ signal decay. Components that fit strongly to a model that assumes temporal dependence and fit poorly with a model that assumes temporal independence are retained. Components were identified and categorized using default options in AFNI’s tedana.py function. Subsequently, data from each scan run were aligned and registered to each participant’s T1 image. Data from each participant were then resampled into 3-mm isotropic voxels and spatially transformed into Talairach atlas space (Talairach and Tournoux, 1988).

### Quantification and Statistical Analysis

#### fMRI Task Analyses

Functional scans consisted of 237 MR frames (233 after initial frames were discarded), with all task scans lasting 8 minutes 42 s in duration. Two scan runs were used from each participant. Overall motion summaries for each run were obtained using the AFNI program @1dDiffMag, which estimates the average of first differences in frame-to-frame motion across each scan run and is comparable to mean Framewise Displacement (Power et al., 2012) calculated over a run. All runs from all participants had DiffMag scores less than 0.3 mm/TR, and all were included in the subsequent analyses. Functional data for each participant were normalized voxelwise by the mean signal intensity of each voxel (to % signal change) and detrended using a fourth-order polynomial baseline model, with no additional spatial smoothing applied. Traditional task analysis was conducted using a general linear model (GLM), in which the data at each time point are treated as the sum of all effects thought to be present at that time point, plus an error term. Responses associated with each condition were modeled using TENT functions, with a separate regressor for each time point following the stimulus onset. This approach assumes that all responses for a given condition share the same response shape, but makes no assumption as to what the shape of that response might be. Responses for each participant were modeled over six time points, based on the range of trial durations determined by the stimulus length and inter-stimulus temporal jitter (max of 13.2 s or 6 TRs). One additional regressor of no-interest was coded for error trials (either omissions or commissions). For the purposes of statistical testing, response magnitudes were estimated by averaging the 3^rd^ and 4^th^ time points of the TENT function corresponding to the peak of the typical BOLD response, reflecting activity 4.4 to 8.8s following stimulus onset.

Separate regression analyses were conducted for each run per participant, with beta coefficients calculated in each voxel at 6 time points post-stimulus onset across the correct naming trials. Beta coefficients averaged at the BOLD peak (3rd and 4th time points) were then tested against zero across participants using one-sample t-tests. Voxels with significant positive t-values indicated above-baseline responses (i.e. Positive responses) and voxels with significant negative t-values indicated below-baseline responses (i.e. Negative responses). Voxels with Positive or Negative responses replicating across both runs (p<.05, FDR q<.05) were then identified and clustered spatially into regions of interest (ROIs).

#### fMRI Task-Based Functional Connectivity Analyses

In order to avoid contamination of task-based functional connectivity estimates from the temporal contour of the evoked responses of individual trials, only the average peak BOLD response was retained from each individual trial (the raw average of the 3rd and 4th timepoints of the task residual timeseries following the onset of each stimulus). This resulted in an item series of peak BOLD responses from correct trials in each individual voxel (max of 50 trials per run). Data were further concatenated across runs to maximize statistical power and improve the stability of the connectivity estimates. ROI-average item series were calculated for each of the Negative ROIs for each participant, and these were used in seed-based correlations (Pearson) with all brain voxels across trials. Pearson correlations were Fisher z-transformed to improve normality and then the mean correlation across participants was tested against zero using one-sample t-tests.

While ME-ICA preprocessing does a good job at removing non-BOLD artifacts such as head motion (Power et al., 2018), it fails to remove slower respiration artifacts that can be linked to brief head movements. We therefore developed trial-level nuisance covariate measures of head motion and more omnibus measures of signal amplitude that could reflect the added variance of any unknown artifact signal (tantamount to windowed estimates of temporal signal-to-noise ratio, or tSNR). Motion estimates for each experimental trial were derived by first calculating Framewise Displacement (Power et al., 2012), which provides a TR-based measure of transient head motion. This measure was then averaged over the 1st 4 TRs of each trial, which included the time period of the peak BOLD response, as well as the first two TRs during which most of the motion due to naming would be expected. Two distinct measures of voxelwise signal amplitude were derived, one that calculated the standard deviation of the 1st 4 TRs of each trial relative to the local mean (of the same 4 TRs) and one that calculated the standard deviation of the 1st 4 TRs of each trial relative to the entire run-level mean. For analyses in which nuisance measures were partialled out of correlation estimates, all three measures (trial-level estimates of motion and two estimates of signal amplitude) were removed simultaneously. Partial correlation estimates could then be Fisher z-transformed and tested against zero at the group level using one-sample t-tests. Tests against zero were carried out at both the voxel-level and on the average over all Positive voxels.

#### Beta weight covariation analyses

Voxelwise beta coefficients from the two separate runs were used for these analyses. Each participant contributed a single measure in each voxel, which was the difference of the beta coefficients in run 1 versus run 2. This difference measure was averaged across voxels in the Negative ROIs and was correlated with all of the voxels in the brain across participants. Since each participant’s data was statistically independent of all of the others, the degrees of freedom for this correlation analysis was simply the number of participants (N=40) minus 2. Run-level estimates of motion and mean voxelwise signal amplitude were also partialled from these correlation analyses (the difference of run 1 minus run 2), with the loss of 1 additional degree of freedom per nuisance covariate (1 for motion and 1 for signal amplitude: df = 36; there was no need for two separate estimates of signal amplitude since the mean used was always defined over the entire run). Results were corrected for whole-brain comparisons using FDR to q<.05. Statistical tests were carried out at the voxel level, but descriptive statistics were also tabulated by averaging over all Positive voxels.

#### Correlations with response time

Response times (RTs) calculated in the picture naming task were correlated (Spearman, due to the typical skewing of the RT variable) with the trial-level BOLD response amplitudes on the correct trials in each voxel for each participant. These correlations were then Fisher z-transformed and tested at the group level against zero (either positive or negative mean correlations) using one-sample t-tests. Whole-brain testing was corrected using FDR to p<.05, q<.05.

#### Variance due to positive coupling between Negative ROIs and Positive voxels

Variance in Positive voxels due to positive coupling with Negative ROI trial-level variability was measured using regression and an *R*^*2*^ approach. The trial-level data in each positive voxel served as the dependent variable and the Negative ROI trial-level data served as the independent variable. Variance of the residuals of the regression model was then subtracted from the total trial-level variance in the Positive voxels to determine the *R*^*2*^ of coupling between Negative ROIs and Positive voxels (N->P). *R*^*2*^ of this coupling not due to nuisance variables was determined by calculating the *R*^*2*^ estimates including these variables in the model (independent variables=Negative ROI, motion, two standard deviation measures) and subtracting the *R*^*2*^ estimates when leaving out the Negative ROI predictor variables (i.e. independent variables=motion, two standard deviation measures). The same approach was taken to finding the *R*^*2*^ not due to RT, subtracting the *R*^*2*^ estimate when not including the Negative ROI predictor variable from the *R*^*2*^ estimate of the full model (independent variables=Negative ROI, motion, two standard deviation measures, RT). *R*^*2*^ due to RT in the trial-level Negative ROI data (and Positive voxels) was found by subtracting the *R*^*2*^ estimates when not including RT from the *R*^*2*^ estimates when using the full model (e.g. independent variables=motion, two standard deviation measures, RT; dependent variable=Negative ROI trial-level data).

#### Assessing similarity of seed-based correlation maps and BOLD-RT correlations

Both the seed-based correlation maps from Negative ROIs using the trial-level amplitudes and the run-level beta coefficients were compared to the maps of voxelwise BOLD-RT correlations within the Positive voxel mask. The similarity of these maps was quantified using Spearman rank correlations. However, the appropriate degrees of freedom to assess statistical significance of these correlations are unknown due to spatial autocorrelations in all of these maps (i.e. nearby voxels tend to have similar values). Permutation testing, in which values in the two maps being compared are randomly shuffled across voxels, is not ideal either as this procedure does not result in random data that are matched in spatial autocorrelation to the original maps. We therefore conducted Monte Carlo simulations using random data with spatial autocorrelations matched to the original data using the AFNI programs 3dFWHMx and 3dClustSim, which empirically determine the spatial autocorrelation function of the data. 10,000 matched random maps were correlated (Spearman rank) with the actual BOLD-RT correlation map in order to derive a null distribution, against which the observed correlations of the actual data were compared. Two-tailed p-values were then calculated from the rank order of the actual data relative to the null distribution, with FDR correction applied for multiple comparisons (p<.05, q<.05). Negative values of the observed correlations between maps would be consistent with competition plus a non-specific global factor, whereas positive correlations would be consistent instead with cooperation.

### Data and Code Availability

The data supporting the current study have not been deposited in a public repository because subsequent analyses are ongoing. Data are available from the corresponding author on request.

## Supplemental Information Titles and Legends

**Table S1. Positive and Negative regions**. Negative ROIs and associated centers of mass (Talairach-Tournoux coordinates) and sizes. Related to Figure 2.

**Table S2. Prediction 1 results: seed-based correlation analyses**. Average correlation (*r*) values over all Positive voxels for prediction 1 analyses, with associated standard errors of the mean (SE). Beta coefficient covariation analyses were calculated only at the group level and lack standard error estimates. Related to Figure 3.

**Table S3. Prediction 2 results: voxelwise correlation between trial-level BOLD magnitude and RT**. Average correlation (r) values, standard errors of the means (SE) and *p*-values (*p*) for prediction 2. Note: *p*-values for Positive voxels showing a negative BOLD-RT correlation were not included as these voxels were identified on the basis of displaying a significantly negative BOLD-RT correlation. Related to Figure 4.

