## Supplementary Materials for "Brain regions with positive and negative task-evoked responses engage in cooperative rather than competitive interactions"

**Table S1. Positive and Negative regions.** Negative ROIs and associated centers of mass (Talairach-Tournoux coordinates) and sizes. Related to Figure 2.

| **ROI** | **Center of Mass**  **(Left-Posterior-Inferior)** | **Voxels (3 mm^3^)** |
| --- | --- | --- |
| Positive Mask | | |
| All Positive Voxels | X = -4.5, Y = -34.5, Z = -42.5 | 26,114 |
| Negative ROIs | | |
| 1 Posterior Cingulate Cortex | X = +1.3, Y = -55.0, Z = +28.7 | 241 |
| 2 Right Angular Gyrus | X = +47.2, Y = -59.7, Z = +31.7 | 202 |
| 3 Ventromedial Prefrontal | X = +2.5, Y = +49.0, Z = +3.6 | 198 |
| 4 Right Superior Frontal Gyrus | X = +24.8, Y = +20.8, Z = +46.5 | 99 |
| 5 Left Angular Gyrus | X = -45.3, Y = -60.8, Z = +33.0 | 86 |
| 6 Right Anterior STS | X = +56.8, Y = -17.4, Z = -14.3 | 69 |
| 7 Right Frontal Pole | X = +25.8, Y = +57.5, Z = +13.4 | 58 |
| 8 Right Middle Frontal Gyrus | X = +39.2, Y = +12.0, Z = +42.4 | 14 |

**Table S2. Prediction 1 results: seed-based correlation analyses.** Average correlation (*r*) values over all Positive voxels for prediction 1 analyses, with associated standard errors of the mean (SE). Beta coefficient covariation analyses were calculated only at the group level and lack standard error estimates. Related to Figure 3.

|  | Seed-Based Correlation | | Seed-Based Correlation, Partialling Nuisance Covariates | | Beta coefficient Covariation Analysis, Partialling Nuisance Covariates (df 39) |
| --- | --- | --- | --- | --- | --- |
| Negative  ROI | r | SE | r | SE | r |
| 1 Posterior Cingulate Cortex | 0.261 | 0.018 | 0.245 | 0.017 | 0.289 |
| 2 Right Angular Gyrus | 0.237 | 0.019 | 0.221 | 0.018 | 0.238 |
| 3 Ventromedial Prefrontal | 0.258 | 0.018 | 0.239 | 0.016 | 0.215 |
| 4 Right Superior Frontal Gyrus | 0.286 | 0.017 | 0.270 | 0.017 | 0.285 |
| 5 Left Angular Gyrus | 0.241 | 0.018 | 0.224 | 0.017 | 0.266 |
| 6 Right Anterior STS | 0.212 | 0.020 | 0.195 | 0.018 | 0.223 |
| 7 Right Frontal Pole | 0.260 | 0.017 | 0.242 | 0.016 | 0.210 |
| 8 Right Middle Frontal Gyrus | 0.248 | 0.017 | 0.231 | 0.017 | 0.170 |

**Table S3. Prediction 2 results: voxelwise correlation between trial-level BOLD magnitude and RT.** Average correlation (r) values, standard errors of the means (SE) and *p*-values (*p*) for prediction 2. Note: *p*-values for Positive voxels showing a negative BOLD-RT correlation were not included as these voxels were identified on the basis of displaying a significantly negative BOLD-RT correlation. Related to Figure 4.

| ROI | r | SE | *p* |
| --- | --- | --- | --- |
| Positive Mask | | | |
| All Positive Voxels | 0.033 | 0.008 | 1.21e^-4^ |
| Positive Voxels Showing Negative BOLD-RT Correlation | | | |
| Left Supramarginal Gyrus | -0.0647 | 0.0181 | - |
| Right Supramarginal Gyrus | -0.0745 | 0.0152 | - |
| Left Inferior Frontal Gyrus | -0.0618 | 0.0135 | - |
| Right Inferior Frontal Gyrus | -0.0584 | 0.0118 | - |
| Right Putamen (anterior) | -0.0486 | 0.0113 | - |
| Right Putamen (posterior) | -0.0533 | 0.0117 | - |
| Negative ROIs | | | |
| 1 Posterior Cingulate Cortex | -0.074 | 0.017 | 7.39e^-5^ |
| 2 Right Angular Gyrus | -0.036 | 0.013 | 0.0114 |
| 3 Ventromedial Prefrontal | -0.049 | 0.014 | 0.0013 |
| 4 Right Superior Frontal Gyrus | -0.038 | 0.012 | 0.0044 |
| 5 Left Angular Gyrus | -0.043 | 0.017 | 0.0147 |
| 6 Right Anterior STS | -0.025 | 0.010 | 0.0133 |
| 7 Right Frontal Pole | 8.16e^-5^ | 0.011 | 0.994 |
| 8 Right Middle Frontal Gyrus | 4.43e^-4^ | 0.014 | 0.975 |
